# Assessing gait control from short time series using a geometric characterization of stride-by-stride corrections

**DOI:** 10.64898/2026.11.26.769496

**Authors:** Clémence Vandamme, Renaud Ronsse, Frédéric Crevecoeur

## Abstract

Gait variability in healthy adults exhibits Long-Range Autocorrelations (LRA), reflecting long-term statistical dependencies between stride intervals. Previous studies have interpreted LRA within an optimal control framework, where stride length and duration are regulated in a coordinated manner to maintain a constant walking speed. Such objective offers redundancy among gait parameters that achieve success. In this context, imposing constraints on the walking cadence reduces this redundancy and leads to a loss of LRA. The model therefore predicts that during free walking, stride-by-stride adjustments should be directed toward the redundant axis, whereas during cued walking, they should be directed toward the imposed target values. Consequently, differences between conditions should be observable based on a geometric characterization of changes in length and duration. With simulations and experimental data, we show that stride-by-stride adjustments discriminate between cued and free walking conditions, supporting the geometric interpretation of the model. Moreover, this metric achieves good discrimination (Area Under the Curve = 0.74) for 50-stride series, compared with LRA estimates that typically require series of 80 strides to achieve comparable success. In sum, our study validates a novel geometric approach to quantifying gait control strategies, supporting the emergence of LRA through flexible adjustments of stride parameters.

**Author summary:** While Walking, we constantly adjust the length and timing of each step to maintain a stable walking speed. These adjustments are not entirely random: they exhibit patterns that persist over time, known as long-range autocorrelations. Such patterns have been associated with the ability to adapt movements to changing demands, and their reduction has been linked to an increased risk of falls in older adults and people with neurological conditions. However, measuring these patterns reliably requires long stride series, limiting their practical use. In this study, we developed a new approach to characterize how people regulate their walking by examining how step length and timing change together from one stride to the next. We demonstrated that this approach reliably discriminates between free and cued walking and remained effective with recordings as short as 50 strides. Beyond reducing the amount of data required, our approach provides a more direct description of the strategies used to regulate walking. Therefore, this new metric represents a promising complementary approach for investigating changes in gait control. This approach may also help pave the way for future applications in clinical populations, where shorter walking recordings are particularly valuable.

## Introduction

Gait variability exhibits Long-Range Autocorrelations (LRA), meaning that the autocorrelation function of fluctuations across walking steps is characterized by a slow decay [1]. Mathematically, stride intervals observed at a given moment remain statistically dependent to those that occurred much earlier in time through a power-law dependency. This decay is quantified by a scaling exponent, that can be estimated using different methods, such as Detrended Fluctuation Analysis (DFA) [2] or Adaptive Fractal Analysis (AFA) [3]. In gait, LRA have been associated with flexibility and adaptability, whereas their reduction or disappearance has been linked to an increased risk of falls in older adults and in some pathologies such as Parkinson’s disease [4–6]. There is therefore a significant fundamental and clinical interest in identifying the mechanisms underlying the emergence of LRA in human gait.

Previous work has interpreted LRA within an optimal control framework, in which stride length and stride duration are regulated in a coordinated manner, to preserve a stable walking speed [7]. Because an infinite amount of combinations of these parameters can achieve the same objective, the task objective possesses a degree of redundancy. The absence of systematic stride-to-stride corrections in either duration or length promotes persistent fluctuations and contributes to the emergence of LRA. Conversely, applying stricter control toward one specific combination of stride length and duration reduces persistence leading to more random patterns. Similarly, when external constraints are introduced, such as auditory cues through a metronome, redundancy is reduced, inducing an anti-persistent dynamics [8]. In an adaptation of the model presented in Dingwell et al. (2010) [7], we showed that regulation of the integral of strides interval, which can be seen as synchronization phase errors, reproduced this anti-persistent dynamics [9].

Based on this model, we reasoned that a geometric characterization of stride-by-stride adjustments in a phase diagram representing step length and cadence should in principle capture the differences in gait control strategies corresponding to conditions with or without cues. More specifically, the 2-dimensional vector defined by stride-by-stride changes in length and cadence should be oriented either towards the redundant axis corresponding to constant velocity during unconstrained walking, or towards the single point defined by the externally imposed target cadence in the cued condition. Furthermore, as free walking involves the coordinated regulation of stride length and duration, one would expect a relationship between these two variables, whereas cued condition could result in variables that are potentially less correlated.

A clear motivation to develop complementary metric assessing gait control strategies is to derive reliable markers based on relatively short series, as estimating the exponent that characterizes the decay of the autocorrelation function remains a major challenge. Since long-range dependencies are inherently asymptotic, series relatively long are typically required to reliably capture their scaling properties. Several studies, conducted in diverse populations, have recommended using 500 to 600 strides [10–13], although shorter series of about 200 strides may be sufficient when multiple trials are averaged or to detect group differences [14–16]. Improvements of DFA such as evenly spacing or AFA have also been proposed for analyses based on approximately 250 strides [17–19]. In gait experiment, however, acquiring such long series can be difficult, particularly among populations suffering from gait impairments. Moreover, these analyses assume that the system under study remains stationary throughout the observation window, whereas prolonged walking tasks may induce fatigue or reduced attention [20]. This limitation motivates to identify alternative metrics capable of capturing changes in gait control strategies occurring at short time scales without requiring long series.

In this study, we tested two geometric characterizations of stride series, and compared these estimates to a measure of LRA in series of stride durations. In particular, we investigated whether stride-by-stride corrections along the redundant axis, and the correlation between stride duration and length, could discriminate between different control strategies from synthetic data generated by an optimal control model capturing differences between free walking and cued walking conditions [9]. We then compared the results with experimental data and evaluated the robustness across different series length. Simulations and experimental data showed consistent discrimination between cued and free walking, supporting the proposed control models in both condition and validating our geometric approach for characterizing gait control. In particular, measurements of stride-by-stride correction along the redundant axis still showed good discrimination between conditions with only 50 strides whereas comparable discrimination performance with LRA was only achieved with about 80 strides.

## Results

This section is articulated as follows. First we present three regulatory strategies implemented in the model (Fig 1A). Then, as two of these control strategies were designed to reproduce gait patterns observed during treadmill walking with and without a metronome, we compared the simulated gait patterns with experimental data. Participants walked on an instrumented treadmill during four sessions, each consisting of a period of free walking and a period of synchronization to a metronome (Fig 1B). Three metrics were calculated to characterize gait dynamics in both the simulated and experimental data: (1) the AFA exponent, which characterizes the autocorrelation function of series of stride duration; (2) the correlation between stride duration and stride length; and (3) the average stride-by-stride correction along the redundancy axis corresponding to all combinations of step length and cadence associated with a constant velocity (see

**Fig 1.**
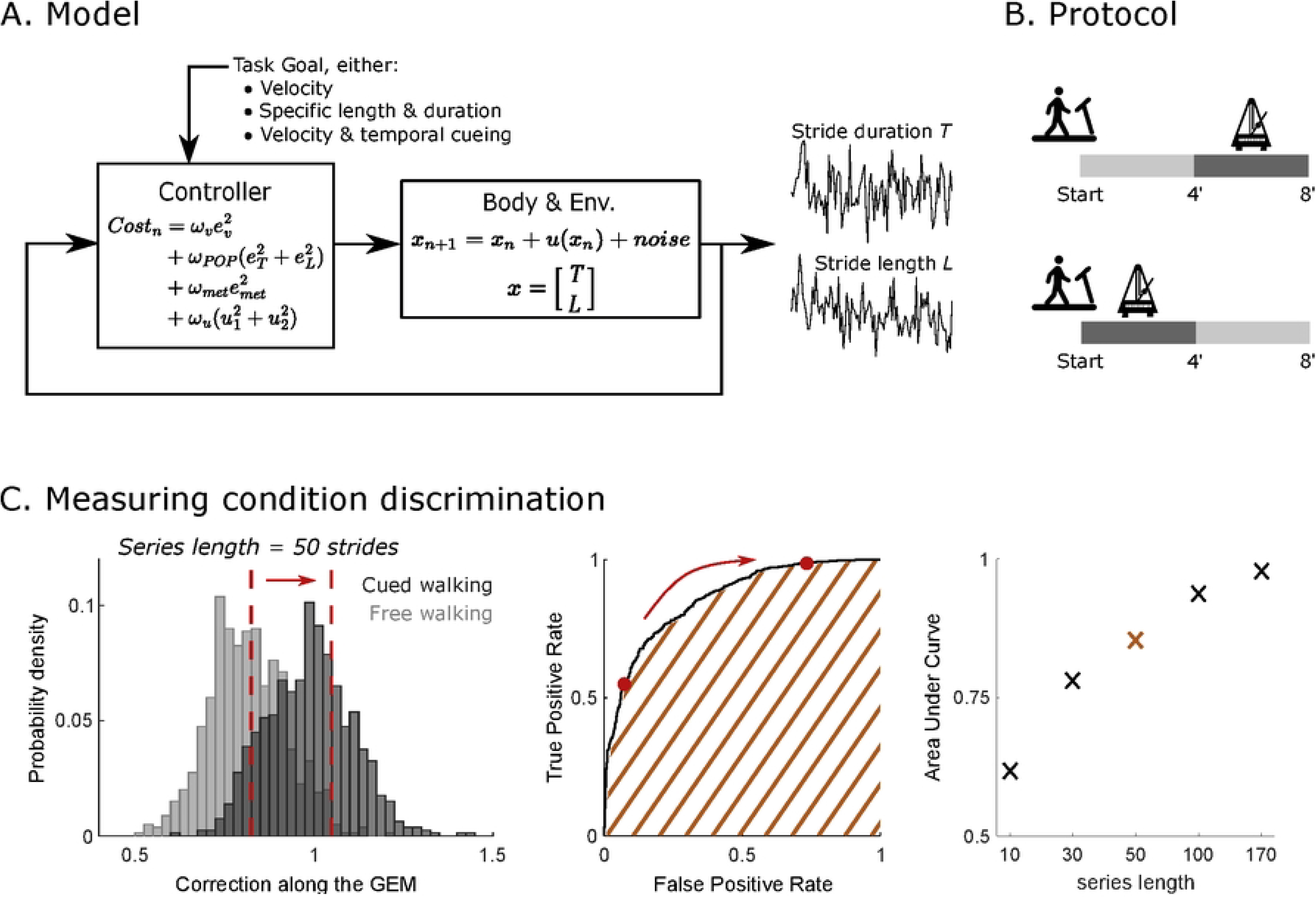
Experimental paradigm, model of stride-to-stride regulation, and procedure for data analysis. A: overview of the two conditions performed by participants on the treadmill, with an illustration of the phases when the metronome was on (dark gray). B: Schematic representation of the model of gait control with three different sets of parameters. C: Illustration of the data processing to measure condition discrimination. Exemplar distribution of the average stride-to-stride correction along the Goal Equivalent Manifold (GEM) for 100 series of 50 strides in cued walking and free walking conditions (left panel). As a threshold (dashed red lines) advances along the x-axis, the true positive rate is represented against the false positive rate (middle panel) with the three red points corresponding to the red lines in the left panel. The area under the curve (AUC) is reported in the right panel for different series lengths. The same procedure was applied to both other estimates (i.e., the AFA exponent and the correlation between stride length and duration) and to both simulated data and experimental data.

Methods). We then evaluated the ability of these estimates to discriminate between conditions using standard classification techniques (Fig 1C), and compared discrimination across series of different lengths. Finally, as participants experienced abrupt transitions between both conditions, initial asynchronies with the metronome could occur and substantially influence gait parameters. Our approach could be leveraged to capture and characterize the effects of this initial asynchrony. The last section provides a full treatment of this aspect.

### Model description

A representative series of stride duration for each strategy is shown in Fig 2A-C. Schematic representations of each control mode, indicated by black arrows, together with an exemplar stride distribution are also illustrated in Fig 2D-F. Three dimensions could be regulated: the stride amplitude, the stride duration and the synchronization with the metronome. Thus, we selected the most relevant variables for each condition: stride length versus stride duration (left and middle panels) and synchronization phase versus stride duration (right panel).

**Fig 2.**
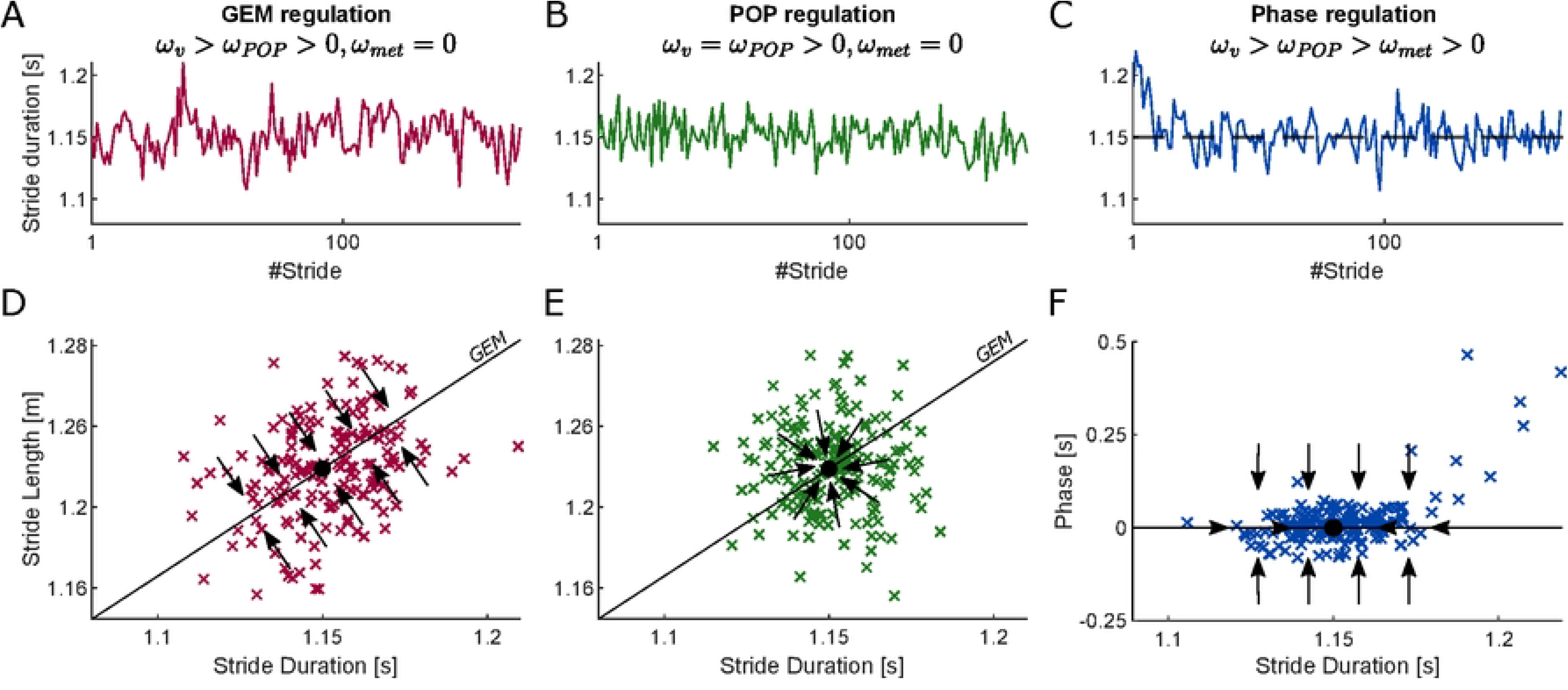
**Exemplar simulated stride series**. A-C: Simulated series of stride duration generated using three different regulatory strategies. D-F: Schematic representation of the control strategies (black arrows) in the relevant state spaces, along with the resulting stride distributions. In D and E, stride length is plotted against stride duration; the black line denotes the Goal Equivalent Manifold (GEM), and the black dot indicates the Preferred Operating Point (POP). In F, synchronization phase is plotted against stride duration; the black line denotes perfect synchronization, and the black dot represents the metronome period (i.e., the interval between two consecutive beats).

The first condition corresponds to unconstrained treadmill walking, as previously described by Dingwell and colleagues [7]. The primary task being to maintain a constant speed, determined by the belt speed, strides are corrected towards a goal-equivalent manifold (GEM), representing all pairs of stride duration (*T* ) and length (*L*) that produce the same walking velocity. The absence of systematic correction towards a fixed target duration results in series exhibiting LRA (Fig 2A, red, AFA *>* 0.5). In the (*T* , *L*) plane, stride parameters display an elliptical distribution with the principal axis aligned with the GEM. In the second regulatory mode, a stronger regulation toward the Preferred Operating Point (POP) was induced, reducing the task redundancy. The series of stride duration gets closer to random noise and the distribution in the (*T* , *L*) plane becomes approximately circular around the POP. While this mode of regulation imposes strict regulation of the walking frequency, it does not guarantee that the timing of the heel strike corresponds to the beat of the metronome. Indeed, the stride duration could be equal to the imposed interval, but out of phase with the metronome. This requires an additional cost parameter, as described in Vandamme et al. (2025) [9], in order to capture synchronization to isochronous cues. Series displayed anti-persistent dynamics, with a small transient trend at the beginning of the series (Fig 2C), reflecting the correction of the initial asynchrony with the rhythmic cueing. In Fig 2F, we observe a tight distribution around the line representing perfect synchronization (phase = 0), which was the primary objective. To keep strides synchronized with the metronome, it was necessary to aim for a specific duration that matches the metronome’s pace. When the stride and metronome are out of phase, however, the target duration must be adjusted, as suggested by the few points scattered away from the synchronization line. This regulation still occurred along the GEM to maintain a constant velocity, but without a tendency towards a specific point, since the target duration depended on the phase error. We observed that such initial and transient phase correction had a major impact on estimates of AFA. Therefore, in the analyses that follow, the first 10 strides of each series generated using phase regulation were removed to avoid confounding effects caused by initial asynchrony. Similarly, the first 10 strides of the experimental series in cued condition were excluded. A fuller treatment of this effect will be described in subsequent analyses.

To characterize these three modes of regulation, we computed three indices characterizing the autocorrelation function or the geometric properties of the distributions: (1) the AFA exponent of the series of stride duration as estimate of slow decay in the autocorrelation function, (2) the correlation between stride duration and stride length to measure the dispersion along the GEM, and (3) the average stride-to-stride correction along the GEM. Fig 3A displays the distribution of these metrics for 100 series of 170 strides from each regulation mode. In line with previous research, POP regulation (green) and phase regulation (blue) resulted in series exhibiting similar AFA exponents, whereas GEM regulation (red) produced higher exponents (left panel). In the middle panel, we observed that the GEM regulation resulted in positive correlations between stride durations and stride lengths, as hinted by the elliptical distribution shown in Fig 2B. Similarly, phase regulation displayed positive correlations, with lower values (Fig 3A, middle panel, blue). In contrast, no correlation on average was observed with the POP regulation. Finally, the correction along the GEM (right panel) separated the three modes more clearly, with the largest corrections observed for POP regulation, intermediate values for phase regulation, and the smallest corrections for GEM regulation.

**Fig 3.**
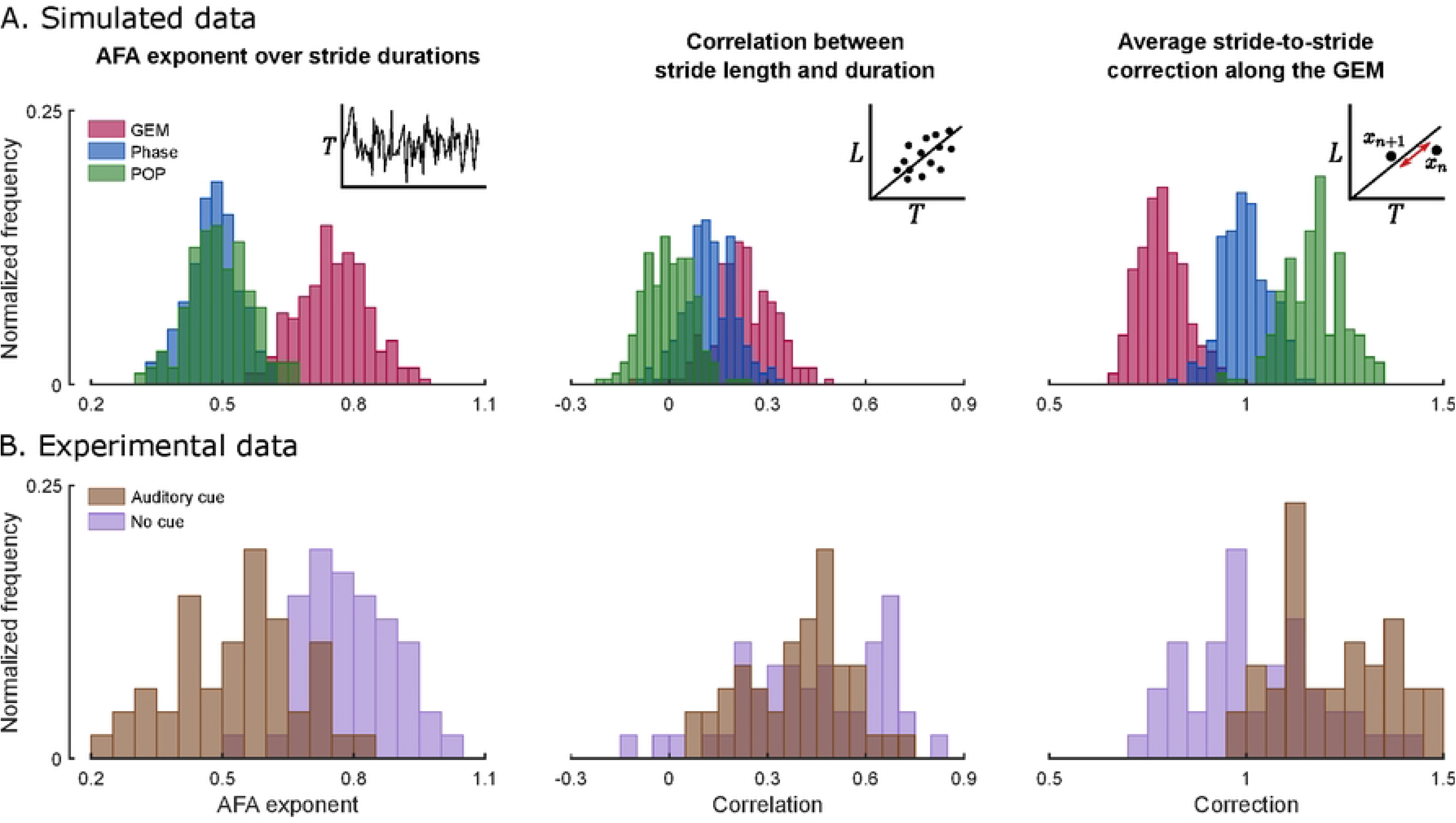
Distribution of three estimates characterizing the strides series. A: Distributions of the AFA exponent of series of stride durations (left), correlation between stride length and stride duration (middle), and the average stride-to-stride correction along the Goal equivalent manifold (GEM) for 100 simulated series of 170 strides in each regulation mode (towards the GEM: red; toward the POP: green; synchronization phase: blue). B: Same metrics computed from experimental data in free-walking (purple) and auditory-cued walking (brown). A total of 94 series of 170 strides, from 15 participants, are represented (47 per condition).

### Comparison with experimental data

In Fig 3B, experimental data were analyzed using the same estimates. Two conditions were tested: a free-walking condition (i.e. without auditory cues), corresponding to GEM regulation [7], and a cued condition (with an isochronous cueing provided by a metronome), which would correspond to phase regulation [9]. Although variability was higher due to inter-individual differences, the results were similar to simulated data. In particular, the distributions of the corrections along the GEM of both conditions displayed much reduced overlap, suggesting that this metric effectively discriminated between both conditions. The same observation applied to AFA exponents, which was expected given that a substantial body of literature had reported markedly different exponents across the two conditions [8, 21, 22]. In contrast, both conditions resulted in similar correlations between stride length and stride durations, which tended to support phase regulation as a candidate model for the cued condition.

While both the AFA exponents and the average corrections along the GEM allowed the two conditions to be distinguished for series of 170 strides, a major limitation of LRA estimation techniques is their limited reliability when applied to short series (e.g. *<* 100 strides). Consequently, in addition to validating the model using metrics other than LRA estimation, a major objective of this study was to identify metrics that remained robust as series length decreases. To this end, we evaluated the ability to discriminate between conditions using series lengths ranging from 10 to 170 strides. The resulting distributions of the average stride-by-stride correction along the GEM are presented in Fig 4A, both for simulated data (top line) and experimental data (bottom line). This analysis focused on the GEM and phase regulation conditions, in line with our primary hypothesis. As expected, the distributions increasingly overlapped as the series length decreased, mainly due to a larger dispersion. To quantify this overlap, we performed receiver operating characteristic (ROC) analyses by plotting the true positive rate against the false positive rate based on a discrimination threshold that varied along the whole domain of each variables (see Methods). The area under the ROC curve (AUC) provided a measure of distributions overlap: a diagonal curve corresponds to complete distributions overlap (*AUC* = 0.5), whereas perfect separation yields a curve reaching the upper-left corner (*AUC* = 1). ROC curves for experimental data (gray) and simulated data (black) are shown in Fig 4B for each series length. Discrimination was generally better in simulations, which was expected given that experimental data included inter-participant variability. In both datasets, ROC curves approached the diagonal as the series length decreased. In the following analyses, a discrimination measure (AUC) was calculated for each participant, then averaged across all participants.

**Fig 4.**
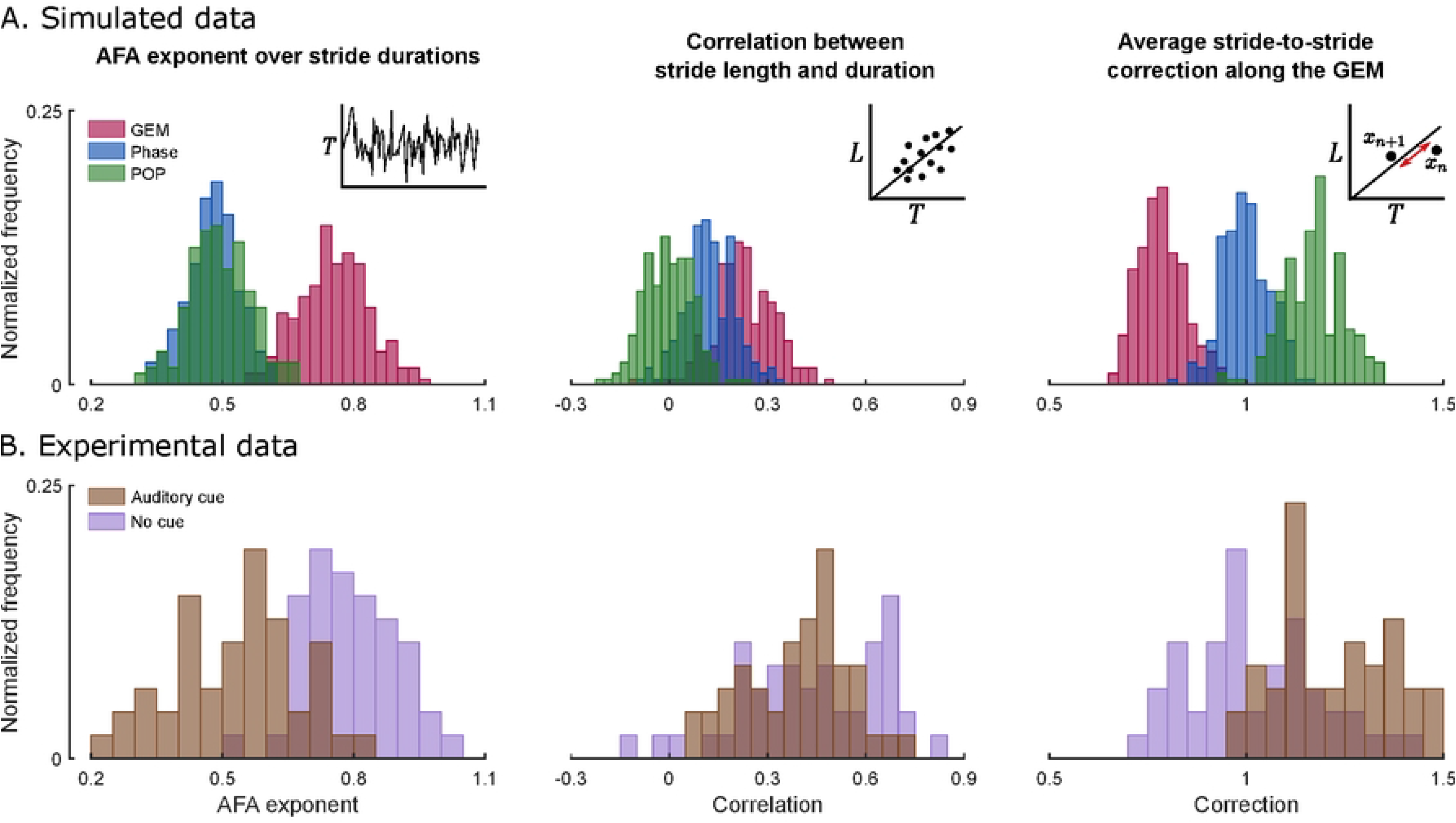
Effect of series length on estimates distributions. A: Distributions of the average stride-to-stride correction along the GEM for increasing series length, from 10 to 170 strides, shown for simulated data (top) and experimental data (bottom). Corresponding Receiver Operating Characteristic (ROC) curve for each series length in simulated (black) and experimental (gray) data.

These individual AUC values of each participant (*n* = 15) are shown in Fig 5 in gray, with the group average indicated by large black dots (dotted line). These values were compared to simulations in two ways: the discrimination between GEM regulation and phase regulation (dashed line, crosses) and the discrimination between GEM regulation and POP regulation (dashed lines, stars). In both case, discrimination was better in simulations for all metrics and series length. However, the experimental results were consistently closer to the simulations involving discrimination between GEM regulation and phase regulation, supporting the hypothesis of a phase regulation in cued walking rather than regulation toward a fixed duration corresponding to the metronome pace. In experimental data, a mixed-effects model revealed a significant effect of series length on the AUC for all metrics (For the AFA exponent: *β*_1_ = 0.003*, t*(58) = 9.8*, p <* 10*^−^*^4^, for the correlation : *β*_1_ = 8 *×* 10*^−^*^4^*, t*(73) = 3.9*, p* = 2 *×* 10*^−^*^4^, for the stride-by-stride correction : *β*_1_ = 0.002*, t*(73) = 14.05*, p <* 10*^−^*^4^). The AFA exponent showed high AUC for long series but decreased rapidly as the series length shortened, especially in experimental data. The correlation between stride duration and stride length provided lower, or even no, discriminability between conditions, and showed substantial inter-participant variability. In contrast, the correction along the GEM yielded high discriminability in both simulated and experimental datasets, with only two participants showing imperfect separation in series of 170 strides (Fig 5, right panel). Importantly, the decrease in AUC with series shortening was less pronounced for this metric compared with the AFA exponent. In particular, for series of 50 strides, a paired t-test indicated that the AUC obtained from the correction along the GEM was significantly higher than that obtained from the AFA exponent (Δ*AUC* = 0.14 *±* 0.16*, t*(14) = 3.40*, p* = 0.004).

**Fig 5.**
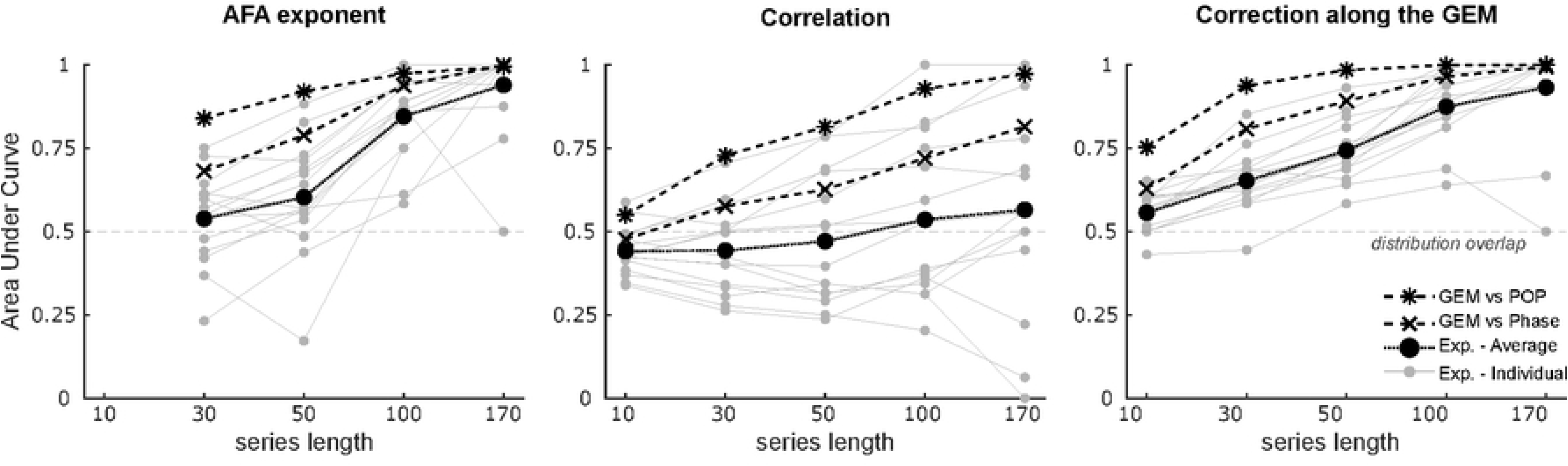
**Discriminability between conditions as a function of series length**. Area under the ROC (AUC) for each participant (gray dots, *n* = 15) and group average (black dots) for the three estimates: AFA exponent (left), correlation (middle) and correction along the GEM (right). Experimental data are compared to simulations for discrimination between GEM and phase regulation (crosses) and between GEM and POP regulation (stars).

### Impact of initial asynchrony in cued condition

In this section, we focused on the effect of the initial asynchrony in the cued condition (for experimental data) and in the phase regulation (for simulated data). In a previous study, we reported that this parameter could have a significant impact on the AFA exponent at the transition between free walking and synchronization with a metronome [9]. A non-zero initial asynchrony was indeed necessary to reproduce a transient increase in the evolution of the AFA exponent at the transition. Here, participants also experienced transitions between free and cued walking. This could affect the AFA exponents, as previously reported, and thus the differentiation between conditions. Furthermore, the initial asynchrony could also impact the correlation between stride length and stride duration, and the average correction along the GEM.

We first examined the effect of the initial asynchrony on the three metrics characterizing stride series in the simulated data. The initial synchronization error in the phase regulation condition was randomly drawn from a uniform distribution in the interval 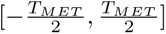. Fig 6A shows the AFA exponent, correlation, and GEM correction as a function of this parameter (blue dots, one per series), together with the average values obtained in the free condition. In the left panel, we observed that larger initial phase error produced higher AFA exponents in the cued conditions, closer to those observed in the free condition, consistent with [9]. A similar pattern was observed for the two other metrics, i.e., the correlation between stride length and stride duration and the average correction along the GEM. This was explained in the model by the fact that larger initial synchronization errors involve transient regulation that deviated from the steady-state duration, thereby promoting greater excursion along the GEM while asynchrony was reduced. Therefore, higher initial asynchronies yielded values closer to free walking conditions, reducing discrimination between conditions.

**Fig 6.**
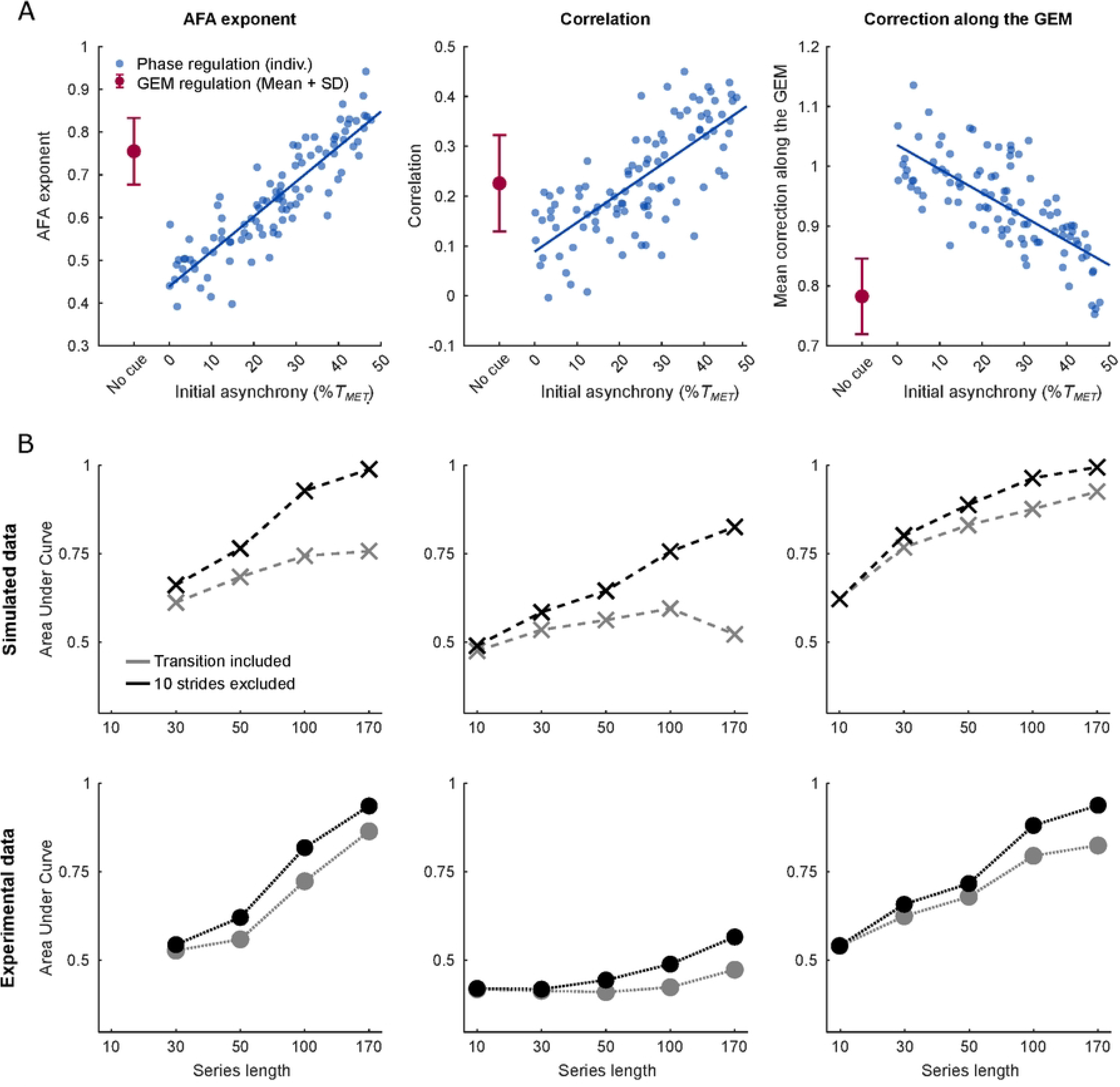
**Impact of initial synchronization errors**. A: Simulated AFA exponent (left), correlation (middle), and GEM correction (right) as a function of initial synchronization error in the phase regulation condition (blue dots, *n* = 100), compared with average values from the GEM regulation. The error bar represents the standard deviation (*n* = 100). B: Comparison of AUC values computed with (gray) and without (black) the initial transition period (10 strides) in simulated data (upper panels, crosses) and experimental data (lower panels, dots)

To verify this effect, we compared the AUC obtained after removing the 10 first strides of the cued condition, with the AUC computed when the transition period was retained, both in simulated and experimental data. As shown in Fig 6B, the discrimination was indeed better when excluding the transition in both simulated data (top panel) and experimental data (lower panel). We also observed a larger influence of this exclusion for series of 170 strides, because shorter series are subsets of the 170-stride series. Therefore, part of them already contained no initial asynchrony prior to exclusion of the transition. In simulations, removing more than 10 strides (e.g., 30 strides) did not further improve discrimination (data not shown), suggesting that removing 10 strides was sufficient to eliminate the biases caused by the asynchrony. In the experimental data, we also compared the two transition directions: from cued to free walking and from free to cued walking. For the former, the recording started once the participant had successfully synchronized with the rhythmic cues, thus implying zero initial asynchrony. In contrast, during the cued portion of the latter condition, the time series could contain transient non-stationarities resulting from the initial synchronization error. As expected, discrimination was consistently higher for transitions from cued to free walking across all metrics, with the strongest effect observed for the correlation.

### Results summary

Overall, we validated a novel geometric approach supporting the hypothesis that LRA emerged from a flexible regulation of gait parameters. In particular, stride-by-stride correction along the GEM emerged as a robust indicator for gait control, maintaining discrimination between conditions for shorter time series. In addition, across simulated and experimental data, free walking was consistently best characterized by regulation along the goal-equivalent manifold (GEM), whereas cued walking was more accurately described by a phase regulation mechanism rather than by a strict convergence toward a fixed preferred duration. While previous work had validated this framework through the presence of (anti-)persistence, the present findings extend this validation by demonstrating that two additional metric independent of the time structure, but guided by the interpretation of the theoretical model, also lead to consistent conclusions. Finally, our model allows a detailed description of the effect of initial asynchrony. These effects had consequences on discrimination between conditions visible in the experimental data, that closely mirrored the ones found in simulated data.

## Discussion

The present study builds upon the interpretation that LRA naturally emerges during free walking from the selective regulation of stride duration and stride length to maintain a stable walking velocity [7]. Recently, we extended this framework to cued walking by proposing that synchronization with a metronome relies on the regulation of the integral of stride durations, effectively compensating for accumulated synchronization errors [9]. Both models were previously validated by reproducing the persistence and anti-persistence observed in stride duration series. The present study extended this validation by demonstrating that two additional metrics, independent of the autocorrelation structure, led to consistent conclusions regarding the underlying control strategies. Using simulations and experimental data, we showed that a geometric characterization of stride-by-stride correction successfully distinguished between free and cued walking, reproduced the effects previously observed on LRA, and captured the influence of initial synchronization errors during transitions to auditory cueing. Importantly, this indicator remained effective for substantially shorter stride series. Together, these findings provided additional validation of the proposed control framework and demonstrate that geometric indicators of gait regulation can offer powerful complementary information about gait control.

Notably, the experimental data in cued condition were more consistent with the phase-regulation model than the POP-regulation. As these two strategies yielded similar AFA exponent, other factors were required to formally determine which one was used in cued walking. The distinction between these two strategies was fundamental. Under POP regulation, each stride were corrected toward a predefined stride duration, independently of previous strides. In contrast, phase regulation explicitly incorporated the current phase error relative to the metronome. Consequently, corrections depended not only on the present stride duration but also on preceding timing errors, often requiring temporary over-corrections to restore synchronization.

In this study, two main elements allowed us to distinguish those two modes of regulation. First, the presence of correlations between stride length and stride duration as observed experimentally and in the model when the phase error was regulated provided an argument against a strict POP-based strategy. Under POP regulation, corrections converged toward a fixed target, producing a circular distribution of strides around this target in the stride duration–stride length plane. This specific distribution was evidenced by the absence of any correlation between these two gait parameters in the simulations, which was not compatible with the correlations observed in the experimental data (Fig 3). Under phase regulation, the target stride durations changed at each stride and gait remained constrained by the need to maintain treadmill velocity. Changes in duration were therefore accompanied by corresponding changes in stride length, preserving a positive correlation between the two variables.

Second, phase regulation introduced an additional variable, namely the asynchrony, that was absent in the other regulation modes. Depending on its initial value, a transitional period of several strides might be present within the series (Fig 2C), skewing the values of the metrics used to characterize the complete series. Removing this trend from simulated data in phase regulation improved discrimination between conditions for all metrics. This effect was well reproduced in the data, as removing 10 strides from the beginning of each series improved the discrimination between conditions for all metrics.

The present study was conducted during treadmill walking, where maintaining a constant velocity was an explicit task constraint, resulting in a well-defined GEM relating stride duration and stride length. Although previous studies have shown that treadmill walking does not alter the temporal structure of stride duration variability in healthy individuals [23, 24], the extent to which our findings generalize to overground walking remains unclear. Overground walking allows velocity to fluctuate naturally and may therefore increase variability within the state space. Consequently, the geometric characterization described here would likely become less sharply defined, which could result in greater overlap between conditions and reduced discriminability. Future studies should therefore investigate whether the proposed geometric framework can be extended to overground walking, where accounting for natural fluctuations in walking speed may be necessary to adequately characterize stride-by-stride regulation.

A major motivation for this work was to identify markers of gait control that remain informative when only short stride series are available. Because LRA are fundamentally defined by their asymptotic behavior, accurate estimation of scaling exponents typically requires long recordings, often in the range of 500-600 strides [10, 11, 13]. Whereas AFA estimated a scaling property across multiple temporal scales [19], the correction along the GEM was computed from consecutive stride pairs and can thus be estimated from a much smaller number of observations. From a methodological perspective, we therefore expected such measures to be less sensitive to series length. The present results supported this prediction. Although the AFA exponent successfully discriminated between free and cued walking for long recordings, its discriminative power decreased rapidly as series length was reduced. In contrast, the average stride-by-stride correction along the GEM maintained substantially higher discrimination, remaining effective even for series as short as 50 strides (AUC = 0.74).

Importantly, our analyses focused on the ability of the metrics to discriminate between regulatory strategies rather than on the estimation of the scaling exponent itself. Previous studies have shown that DFA might still detect group differences with recordings shorter than those generally recommended for reliable exponent estimation. In that sense, series of 100-200 strides could be sufficient to detect difference of 0.1 in DFA exponent, provided that the sample size was large enough [15]. Likewise, stitching together five trials of 25 strides into longer series enabled discrimination between people with Parkinson’s disease and healthy controls using both AFA and DFA [16]. Our results suggested that geometric measures may provide a similarly useful alternative for distinguishing gait control strategies in short recordings. Furthermore, a strong correspondence was observed between AFA exponent and average stride-by-stride corrections along the GEM across both simulated and experimental datasets. Although this has not been specifically tested here, geometric metrics could therefore have a predictive value for estimating scaling exponents in a statistical sense.

This robustness to shorter series is particularly relevant for clinical contexts in which collecting hundreds of consecutive strides may be impractical. Patients with gait impairments, or older adults, may be unable to perform sufficiently long walking trials, limiting the applicability of traditional LRA analyses. However, these analyses are clinically relevant, as reduced LRA have frequently been reported in aging and neurological disorders, such as Parkinson’s disease, and have been interpreted as reflecting impaired flexibility and adaptability of gait control [4, 6, 25]. Within the framework proposed here, it was proposed that such reductions could arise from stronger corrections toward specific operating points and a diminished exploitation of redundancy, similarly to the POP-regulation [7, 26]. Although pathological populations were not investigated in the present study, the model makes explicit predictions that could be tested in future work. If the apparent reduced persistence is due to an increased regulation toward a fixed gait parameter, geometric measures such as the correction along the GEM should differ from those observed in healthy participants. Because these metrics remained informative for relatively short recordings, they might prove particularly valuable in clinical populations for whom long walking trials are difficult to obtain. Beyond their potential robustness to shorter series, it may also offer a more direct and interpretable description of gait regulation. Whereas a scaling exponent provides a global description of temporal structure, geometric metrics directly characterize the regulatory mechanisms that are hypothesized to generate persistence properties.

In conclusion, the present study supports the view that LRA emerge from flexible regulation within a redundant system. By combining simulations and experimental data, we demonstrated that geometric characterization of stride-by-stride corrections provided a meaningful, complementary description of gait control. In particular, correction along the goal-equivalent manifold emerged as a reliable marker capable of discriminating between regulatory strategies even in short recordings. These findings not only strengthen the interpretation of LRA in gait variability but also open new perspectives for the assessment of gait regulation in both experimental and clinical settings.

## Materials and methods

### Experimental data

Fifteen healthy young adults (7 females, age = 29.3 *±* 5.9 years, body mass = 77.4 *±* 17.0 kg, height = 1.75 *±* 0.10 m, leg length = 0.92 *±* 0.05 m) were enrolled in the study between 18/11/2025 and 21/01/2026. Participants were naive to the purpose of the experiment and reported no neurologic, muscular or orthopedic disorder that could alter their gait. The ethics committee of the host institution *(Comité d’éthique hospitalo-facultaire*, UCLouvain, Belgium) approved the experimental procedures and participants provided written informed consent before the experiment following usual procedures.

Participants performed four treadmill walking sessions, each lasting 8 minutes. Sessions consisted of two consecutive 4-minutes periods: one guided with an isochronous auditory cue and one without cueing (Fig 1A). In two sessions, the cued condition was presented first (”Cued*→*Free” sessions), whereas in the other two sessions, the cues were introduced after 4 minutes of free walking (”Free*→*Cued” sessions). Session order was randomized across participants. During all experimental sessions, treadmill speed was held constant.

During cued walking, participants were instructed to synchronize one foot (heel strike) with the metronome. Prior to the four experimental sessions, participants determined their preferred walking speed using the treadmill self-paced mode [27]. Once the preferred speed was established, participants walked for 1 minutes at that fixed speed. The average cadence measured during this familiarization period was used to set the metronome frequency. One participant completed only two sessions, and two participants completed only three sessions, due to time constraints.

Participants walked on an instrumented treadmill (Gaitway 3D, Arsalis, Belgium). Ground reaction forces in three dimensions were measured based on force sensors mounted underneath the treadmill belt. Series of stride duration and stride length were reconstructed by the software. Subsequent analyses and simulations were performed using custom routines implemented in MATLAB R2021a (The MathWorks, Natick, MA). Each series was divided into two segments at the stride corresponding to cue activation or deactivation, occurring at 240 s, to allow separated analyses of the *Cued* and *Free* walking periods. As the number of strides in each series varied across participants due to differences in walking cadence, all series were truncated to 180 strides, corresponding to the shortest available series.

Sessions in which participants failed to reliably synchronize their gait with the auditory cue were excluded from analysis. To identify such sessions, we followed the following procedure : a 50-stride moving average was computed during the metronome phase and the root mean squared error (RMSE) relative to the target metronome cadence was calculated [9]. Sessions were classified as outliers when their RMSE exceeded *Q*3 + 1.5 *× IQR*, where *Q*_3_ denotes the third quartile and *IQR* the inter-quartile range. Based on this criterion, nine sessions over 56 were excluded from further analyses.

### Model

The primary hypothesis was that geometric criteria could discriminate between conditions, based on the model proposed by Dingwell and colleagues [7]. In free walking, gait is regulated towards the Goal Equivalent Manifold (GEM), resulting in series exhibiting long-range statistical persistence. In contrast, stronger regulation toward a single target point is expected to produce more random time series. Within this framework, we extended the model to reproduce cued walking and proposed a formulation in which phase (i.e., the integral of stride durations) is regulated [9]. This section describes these three models. For each of them, 100 time series of 180 strides were generated, matching the series length of the experimental datasets.

In this model, each stride was generated iteratively from the previous one according to the following discrete-time system:

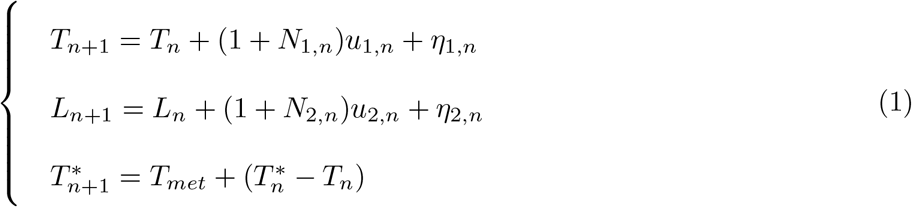

Each stride was characterized by two state variables corresponding to the stride duration *T_n_* and length *L_n_*, with *n* the index of the stride. The noise term consisted in an additive noise vector *η_n_* and a command-dependent noise *N_n_u*. The vector *η* and the diagonal of the matrix *N* were zero-mean, Gaussian random variables with covariance matrix equal to *diag*[*σ*^2^*, σ*^2^], off-diagonal terms being equal to 0. The command *u*(*x_n_*) can be seen as the stride-by-stride regulation of length and cadence against noise disturbances according to the behavioral goal. To incorporate the metronome constraint, a target duration *T^∗^* was introduced. This target represents the duration required to synchronize the next stride with the metronome. It was defined as the sum of the metronome beat interval (fixed and denoted *T_met_*) and the synchronization error from the previous step.

The motor command was determined by a controller to regulate stride variability resulting from internal and external sources of noise according to a behavioral goal. This objective was formalized through a cost function (Eq 2), where each term was weighted by a coefficient *ω* reflecting its relative importance:

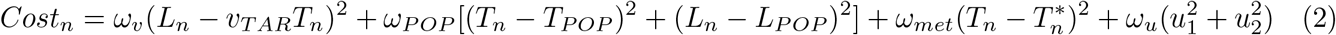

A first objective, weighted by *ω_v_*, was to maintain a constant (self-selected) velocity *v_T AR_*. As an infinite amount of [*T_n_, L_n_*] can produce the same velocity, maintaining constant velocity leaves one degree of freedom in the state variables, making the task redundant. However, along this redundant axis, one pair, referred to as the Preferred Operating Point (POP) and denoted as [*T_P OP_ , L_P OP_* ], may be naturally preferred, likely due to biomechanical constraints on the achievable range of cadence and length modulation. A secondary objective, weighted by *ω_P OP_* , was then to limit deviations from this preferred pair. To penalize phase errors, deviations from the target duration *T^∗^* were additionally penalized with a weight *ω_met_*. Finally, these objectives were balanced against the energetic cost associated with the corrective commands *u*, weighted by *ω_u_*.

To simulate the three different regulatory strategies, we varied the parameters in this cost function as follows. We first set *ω_v_* = 30 and *ω_u_* = 10 in each condition, to account for a strong velocity regulation, required by the treadmill constant velocity. Then, *ω_P OP_* and *ω_Met_* differ from one condition to another. In the GEM regulation, errors are primarily controlled toward the GEM, allowing for some deviations away from the POP. However, to remain within a physiologically realistic range of values, control toward this point is still maintained, to a lesser extent. We set *ω_P OP_* = 2.5 to fit the AFA exponent of series of stride durations obtained in the unconstrained condition. It was also considered that there is no phase control in this case, therefore *ω_met_* = 0. In the POP regulation, we simply increased *ω_P OP_* to 30, giving equal weight to the regulation toward the POP and toward the GEM. Finally, for the phase regulation, we set *ω_met_* = 0.35 to fit the AFA exponent of series of stride durations obtained in the cued condition, similarly to [9]. The penalty over the POP was the same as for the GEM regulation, i.e., *ω_P OP_* = 2.5. All other parameters remained unchanged from one condition to another and are reported in Table 1.

**Table 1.**
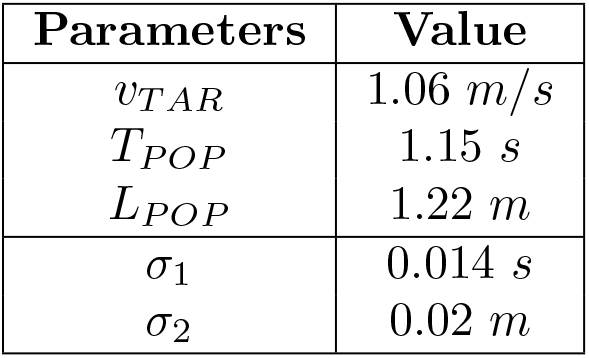
Model parameters

In the phase regulation, we varied the initial asynchrony by manipulating *T^∗^*. The initial value of *T^∗^* could be set to *T_met_*, capturing a perfect synchronization of the first beat of the metronome with the corresponding stride. However, to match the experimental settings more realistically, the initial asynchrony was randomly drawn from a uniform distribution 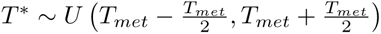. These boundaries were chosen to match the worst initial asynchrony that participants could experience, which corresponds to the case where the metronome beat occurred exactly between two strides.

## Data analysis

To characterize the regulation mechanisms underlying gait dynamics, three metrics were computed from series of stride duration and stride length: (1) AFA exponent of stride duration, (2) the correlation between stride duration and stride length, and (3) the average stride-to-stride correction along the goal-equivalent manifold (GEM). These metrics were applied identically to simulated and experimental datasets.

The **AFA exponent** was used to characterize the temporal structure of series of stride durations. This exponent is linked to the rate of the decay of the auto-correlation function. It is obtained with the Adaptive Fractal Analysis (AFA), a method similar to DFA, described in [3]. Values above 0.5 indicate persistent long-range autocorrelations, values close to 0.5 correspond to uncorrelated fluctuations, and values below 0.5 reflect anti-persistent dynamics.

Next, the **correlation** between stride duration *T* and stride length *L* was calculated using Pearson’s correlation coefficient *r*:

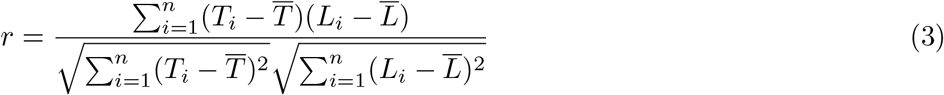

Positive correlations, indicating a relationship between the two variables, were interpreted as reflecting coordinated fluctuations. However, it is important to highlight that the linear regression corresponding to this relationship is not necessarily aligned with the GEM. Correlations close to zero indicated independent regulation of both variables.

Finally, **average stride-by-stride correction along the GEM** was estimated to quantify the magnitude of regulation occurring along the redundant axis. To this end, stride series were first normalized to unit variance. The average walking velocity was then computed as the ratio between mean stride length and the mean stride duration: 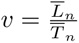 , and the GEM was defined as the line with slope *v*. Next, we computed the absolute distance between each pair of consecutive strides, projected along the GEM, as:

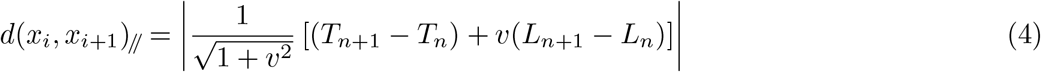

with *i* = 1, 2*, …, n −* 1. The average across all consecutive pairs was then used as the metric of stride-by-stride regulation along the manifold, where larger values reflected higher displacements along the GEM and smaller values indicated less corrections along the manifold.

To compare the regulation strategies, distributions of these metrics were computed from 100 simulated time series generated using each set of parameters and from experimental data obtained during free and cued walking conditions. Based on these distribution, receiver operating characteristics (ROC) analyses were performed to evaluate the ability of each metric to discriminate between regulation modes (Fig 1C). To characterize the overlap between two distributions (left panel), the true positive rate was plotted against the false positive rate while progressively varying the discrimination threshold (middle panel). The area under the ROC curve (AUC) was used as an index of discriminability (right panel), with an AUC of 0.5 indicating chance-level discrimination and an AUC of 1 corresponding to perfect separation between conditions. In experimental data, ROC analyses were performed individually for each participant, and the resulting AUC values were averaged across participants.

These analyses were carried out on series of varying lengths to assess the robustness of the tested metrics when applied to shorter series. In experimental data, the shortest series across participants contained 180 strides. As explained in Experimental data, all series were therefore truncated to 180 strides, discarding strides at the end of the series. To avoid biases linked to the initial asynchrony between heel strike and cues, the first 10 strides of each simulated and experimental cued series were removed from the primary analyses. The same procedure was applied to all series to apply the analyses on series of equal size. The largest series size was therefore 170 strides. Series were then divided into subsets of 100, 50, 30 or 10 strides, with an overlap if necessary, resulting in 2, 4, 6 and 17 subsets per original series, respectively.

Finally, the effect of initial synchronization errors on metrics values and discriminability was investigated in details. To do so, the 10 strides that were initially excluded from the analyses were re-introduced. In addition, 10 strides were removed at the end of the series to obtain series of 170 strides. In simulations, the initial asynchrony was known and uniformly distributed across series between 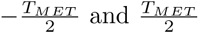. We were then able to evaluate the impact of such asynchrony by recomputing the AFA exponent, the correlation and the average stride-to-stride correction along the GEM for all series. In both experimental and simulations, ROC analyses were repeated with and without removal of the initial transition period. In experimental data, transitions from free to cued walking (Free*→*Cued) were additionally compared with transitions from cued to free walking (Cued*→*Free) to evaluate the impact of synchronization transients on discrimination performance.

### Statistics

We fitted a linear mixed-effects model to evaluate the impact of the series length on the discriminability of each metric, measured with the AUC:

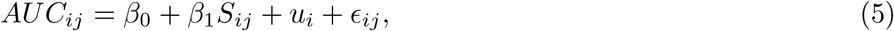

where *S* represents the series length, *i* indexes participants and *j* indexes observations. The term *u_i_* is the random intercept of participant *i* and *ɛ_ij_* the residual error.

In addition, we performed a two-tailed paired t-test to investigate whether the correction along the GEM could better discriminate between conditions than the AFA exponent for short series. We compared the AUC of both metrics for series of 50 strides. The significance level was set at *α* = 0.05.

